# Characterization of pediatric porcine pulmonary valves as a model for tissue engineered heart valves

**DOI:** 10.1101/2023.08.18.553926

**Authors:** Shouka Parvin Nejad, Bahram Mirani, Craig A. Simmons

## Abstract

Heart valve tissue engineering holds the potential to transform the surgical management of congenital heart defects affecting the pediatric pulmonary valve (PV) by offering a viable valve replacement with the compositional, architectural and mechanical properties required to function *in situ*. While aiming to recapitulate the native valve, the minimum requirement for tissue engineered heart valves (TEHVs) has historically been adequate mechanical function at implantation. However, long-term *in situ* functionality of TEHVs remains elusive, suggesting that a closer approximation of the native valve is required. The realization of biomimetic engineered pediatric PV is impeded by insufficient characterization of healthy pediatric tissue. In this study, we comprehensively characterized the planar biaxial tensile behaviour, extracellular matrix (ECM) composition and organization, and valvular interstitial cell (VIC) phenotypes of PVs from piglets to provide benchmarks for TEHVs. The piglet PV possessed an anisotropic and non-linear tension-strain profile from which material constants for a predictive constitutive model were derived. Further, the ECM of the pediatric PV possessed a trilayer organization populated by collagen, glycosaminoglycans, and elastin. Biochemical quantification of ECM proteins normalized to wet weight and DNA content of PV tissue revealed homogenous distribution of proteins across sampled regions of the leaflet. Finally, the predominant phenotype of VICs in the piglet PV was quiescent vimentin-expressing fibroblasts, with a small proportion of activated α-smooth muscle actin-expressing myofibroblasts residing primarily at the base of the leaflet. Overall, the properties characterized in this study can be used to inform TEHV design parameters towards generation of biomimetic pediatric PVs.

**Graphical Abstract:** 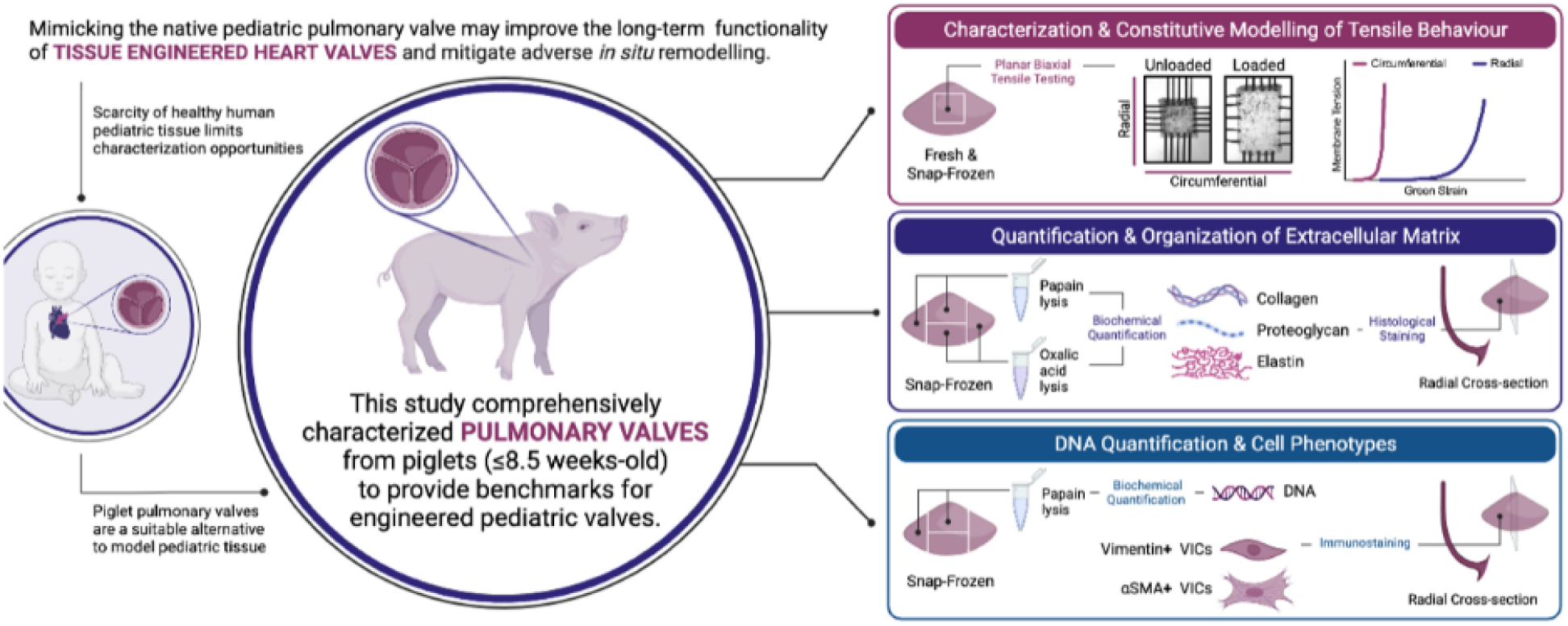

## 1. Introduction

Congenital heart disease (CHD) is the most common congenital disorder affecting approximately 8 in 1,000 live births^1-3^ and the most frequent cause of infant deaths due to birth defects^4^. Approximately 20% of CHDs involve anatomical abnormalities related to the right ventricular outflow tract and pulmonary valve (PV)^5^ that are surgically managed by PV replacement. The demand for pediatric PV replacement is compounded by the increasing frequency of the Ross procedure to treat congenital left ventricular outflow tract obstruction by replacing the aortic valve with a pulmonary autograft^6^. Among available replacements, cryopreserved homografts and bioprosthetic xenograft valves are preferred to mechanical valves for pediatric recipients, due to the restrictions imposed by anticoagulation therapy^7^. However, homografts and bioprosthetic valves are not without their limitations, owing to the limited supply and potential for allosensitization in the former^8^, and the propensity of the latter to undergo structural deterioration and fibrotic stiffening^9^. The limited durability of homografts and bioprosthetic valves necessitates repeated surgical intervention, making the PV the most commonly replaced valve in CHDs^10^.

Heart valve tissue engineering (HVTE) holds the potential to resolve the currently unmet need for a pediatric PV replacement with somatic growth potential and functional adaptation to support life-long performance, obviating the need for repeated intervention. For over two decades, researchers have pursued a range of *in vitro, in situ,* and *in vivo* HVTE strategies, encompassing different progenitor cell sources, biomaterials and mechanical conditioning protocols, with the aim of producing living neotissue with the compositional, architectural and mechanical properties necessary to function *in situ*^11,12^. While the healthy native valve serves as a gold-standard, the minimum requirement for a bioengineered valve at implantation is adequate mechanical function, which is determined by the geometric and material properties of the biomaterial scaffolding and any *de novo* synthesized extracellular matrix (ECM). Experience to date in large animal models indicates that short-term functionality can be achieved for the PV without fully mimicking the native leaflet^12^. However, realization of a bioengineered valve with long-term functionality remains elusive due to universal reports of adverse *in vivo* remodeling, defined by cell-mediated leaflet thickening and retraction. Adverse tissue remodeling is attributable to persistent myofibroblast activation, which can be a function of mechanical stress, inflammation, and/or circulating pro-fibrotic cytokines^13,14^. It is possible that non-physiologic features of tissue engineered heart valves (TEHVs) underly the pro-fibrotic stimuli and an *in vivo*-like structure is required at implantation to provide for long-term function and homeostasis.

There are few studies aimed at comprehensively characterizing the native pediatric PV. This is attributable, in part, to limited access to healthy human pediatric PVs from cadaveric or living donors that are not affected by anatomical abnormalities or systemic disease and trauma. Accordingly, characterization of the pediatric PV are limited to a small sample size spanning donor tissues from early infancy to pre-adolescence^15^. If not limited by a small sample population, characterization of the human pediatric PV has been restricted to histological assessment of the ECM and valvular interstitial cell (VIC) phenotypes^16,17^, providing insight into some but not all relevant properties for HVTE.

In lieu of human pediatric leaflets, TEHVs are often benchmarked against explanted *adult* PVs from large animal models^18-22^ used in preclinical studies of bioengineered constructs. However, evidence of age-related changes in VIC phenotype^16^, ECM composition and organization^15,16,23^, and mechanical behaviour^15,23,24^ emphasize the need to evaluate bioengineered pediatric PVs against their age-matched native equivalent. Pediatric animal models offer an alternative to native human tissue and enable studies on a larger sample population with relevant anatomy and physiology. Due to their analogous cardiovascular anatomy^25^ and systemic hemodynamic variables (e.g., mean blood pressure, heart rate, and cardiac output)^26^, piglet models are often used as a growing heart model for cardiovascular surgical studies^27-29^. However, the mechanical, structural, compositional, and cellular properties of piglet native PVs have not been previously reported.

In this study, we comprehensively characterized the piglet PV as a model for the pediatric human PV. Specifically, we measured the thicknesses and planar biaxial tensile properties of piglet native PV leaflets, determined the content and organization of three constituent structural ECM proteins (collagen, elastin, and glycosaminoglycans), and assessed VIC phenotypes. These data describing native pediatric PV structure and function provide design specifications for pediatric TEHVs and can serve as benchmarks for their assessment.

## 2. Materials and Methods

### 2.1 Pulmonary Valve Isolation and Processing

Hearts from piglets (York-Landrace x Duroc) between 7-8.5 weeks old (55.6 ± 2.5 days old; 14.76 ± 2.74 kg; mean ± SD; N = 10) were provided in-kind by Dr. Robert Friendship (Ontario Veterinary College, University of Guelph). Hearts were transferred in Dulbecco’s phosphate-buffered saline (PBS) (Gibco, USA, cat# 14190-144) (PBS-/- unless otherwise indicated) to the University of Toronto. Within 2 hours of harvest, PVs intact with the pulmonary artery were isolated from the heart and serially rinsed in five PBS baths to remove residual blood from the tissue. PVs were kept hydrated in a PBS bath for up to 4 hours until further processing.

One fresh PV leaflet per valve from 6 donors was isolated and stored in PBS for planar biaxial tensile testing of fresh leaflets as described below. The remaining leaflets and conduit from 6 donors and intact PV from 4 donors were snap-frozen using liquid nitrogen. Snap-frozen PV tissues were stored at -80°C for further analysis. Leaflets from snap-frozen PV tissues were processed for tensile testing, biochemical assays, and immunofluorescent or histological staining. The processing of the PV leaflets for different analyses is summarized in Supplemental Figure 1.

### 2.2 Equibiaxial Force Displacement-Controlled Biaxial Tensile Testing

Tests were performed according to a previously published protocol^30^. A square 4.5×4.5 mm^2^ sample was cut from the belly region of each PV leaflet tissue using a parallel cutter and mounted to a biaxial mechanical tester (BioTester 5000, CellScale, Canada), equipped with 0.5 N load cells, using a tine sample attachment system (Biorakes with 0.7 mm tine spacing, CellScale, Canada; Figure 1A). Graphite powder was sprinkled on the surface of the samples to track the displacement of various points on the tissues during the tests. While submerged in a PBS bath at 37°C, tissues were subjected to ten displacement-controlled preconditioning cycles with displacements of *U*_*Max*,*Circ*._ and *U*_*Max*,*Rad*._ in the circumferential and radial direction, respectively, that generated equibiaxial loading in the tissue. Subsequently, they underwent nine displacement-controlled testing protocols with gradually increasing circumferential-to-radial load ratios (Figure 1B and Supplemental Table S1). The displacement values, *U*_*Max*,*Circ*._, *U*_*Max*,*Rad*._, *U*_*Adj*,*Circ*._, and *U*_*Adj*,*Rad*._ were determined experimentally prior to the test to provide equibiaxial loading in the preconditioning and the fifth test cycles while keeping the maximum load approximately constant across all other test cycles (Figure 1C). All biaxial tensile tests were performed at a strain rate of 0.08 s^-1^. The BioTester software (LabJoy v.10.77) was used to collect force and displacement values in X and Y directions and determine the X-Y coordinates of four points on the samples during the stretch cycles via image tracking.

**Figure 1:**
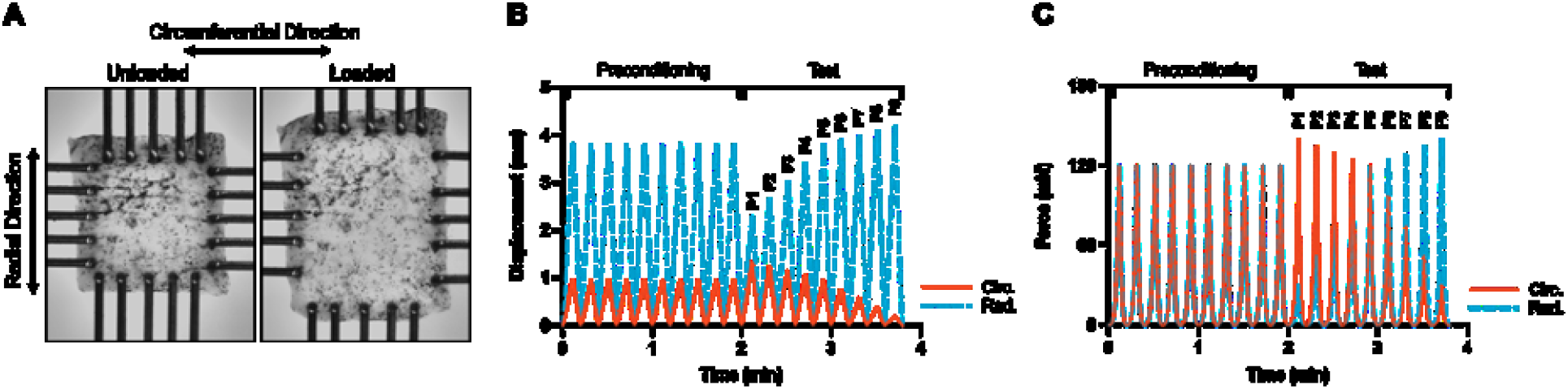
Overview of equibiaxial force planar tensile testing of piglet pulmonary valve (PV). (A) PV tissue sample attached to the Biotester using a tine attachment system, shown in its unloaded and loaded states. (B) An example of prescribed displacement applied in the circumferential (Circ.) and radial (Rad.) direction of a PV tissue sample in the preconditioning and nine test cycle protocols (P1–P9). (C) An example of the resultant force that the PV tissue sample experienced in the preconditioning and test cycles.

Experimental data collected from biaxial tensile tests were used to calculate Green strain, membrane tension, and material constants for a seven-parameter Fung model^31^, as reported in detail previously^30^ (see Supplemental Information). Briefly, assuming that the strain energy is conserved in the test cycles (P1–P9), the tension and Green strain tensors are related according to:

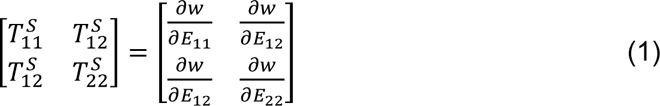

where *w* is the 2-dimensonal strain energy function.

To characterize the mechanical properties of the PV tissue via material constants, a seven-parameter Fung model^31^ was used:

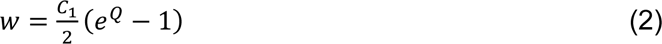

where *Q* = *C*_2_*E*_11_^2^ + *C*_3_*E*_22_^2^ + 2*C*_4_*E*_11_*E*_22_ + *C*_5_*E*_12_^2^+2*C*_6_*E*_11_*E*_12_+2*C*_7_*E*_22_*E*_12_.

All calculations were performed using a MATLAB code previously reported and kindly provided by Dr. M. Labrosse (University of Ottawa)^30^.

### 2.3 Papain and Oxalic Acid Digestion

Snap-frozen PV leaflets were thawed in a water bath at 37°C and leaflets were isolated and kept hydrated in a PBS bath. Leaflet tissue sections from designated regions (edge, belly, periphery) were isolated as described in Supplemental Figure S1. Excess PBS was drawn from the tissue using a Kimwipe™ and samples were weighed prior to tissue digestion and the wet tissue weight of each sample was recorded.

For papain digestion, PV leaflet tissue from each of the three regions were incubated in safe-lock Eppendorf tubes with 1 mL of 40 µg/mL papain (Sigma, cat. #P3125) solution in buffer (35 mM ammonium acetate, 1 mM ethylenediaminetetraacetic acid disodium salt dihydrate, 2mM dithiothreitol). Samples were incubated on a heat-block at 65°C for 72 hours. Papain lysates were stored at -20°C for further analysis.

For digestion in oxalic acid, PV leaflet samples from each region were incubated in safe-lock Eppendorf tubes with 750 μL of 0.25 M oxalic acid. Samples were kept on a heatblock at 100°C for 1 hour and subsequently centrifuged (10,000 x g, 4°C, 10 minutes). The supernatant (oxalic acid lysate) was collected and stored at -20°C for further analysis.

Total DNA, sulfated glycosaminoglycan (s-GAG), and hydroxyproline content of papain digested lysates was quantified using the Hoechst dye 33258 assay^32^, dimethylmethylene blue dye binding assay^33^, and chloramine-T/Ehrlich’s reagent assay^34^, respectively. The insoluble elastin content of oxalic acid digested samples was measured using the Fastin^TM^ Elastin Assay Kit by Biocolor (Accurate Chemical and Scientific Corporation, cat. # CLRF2000). Detailed methodology for biochemical assays are provided in Supplemental Materials.

### 2.4 Immunofluorescent Staining and Imaging

Leaflets embedded in optimal cutting temperature (OCT) compound were sectioned with a cryotome to acquire 7 μm thick radial sections of PV tissue on Fisherbrand™ Superfrost™ Plus Microscope Slides (Thermo Fisher Scientific, cat #22037246). Slides were kept at -80°C prior to tissue staining. To identify quiescent fibroblastic and active myofibroblastic VIC phenotypes, sections were immunostained for vimentin and alpha smooth muscle actin α-SMA. Briefly, slides were thawed at room temperature for 30 minutes. Sections were then fixed with 100% ACS grade acetone (Caledon Laboratories, cat #1200-1) for 2 minutes before air drying for 30 minutes. Slides were washed with PBS and tissue sections were encircled with a PAP pen. After the PAP pen had dried, slides were washed with PBS, and blocked for 10 minutes with 10% (w/v) bovine serum albumin (BSA) (Sigma-Aldrich, cat #A9657) in PBS. The co-stain dye was prepared from a 1:50 dilution of anti-α-SMA CY3 conjugated primary antibody (Sigma, cat #C6198) and 1:50 dilution of anti-vimentin Alexa Fluor™ 488 conjugated primary antibody (Thermo Fisher Scientific, cat #MA5-11883-A488) in 10% BSA. Sections were incubated with the co-stain dye for 1 hour in a dark humid chamber box. Slides were subsequently washed four times for 2 minutes in PBS. Nuclei were then stained with a 1:50 dilution of a 1 mg/mL Hoechst 33324 stock dye (Sigma, cat #B2261) in PBS for 5 minutes. Slides were rinsed twice in PBS for 2 minutes, before washing twice in distilled water for 2 minutes. Stained sections were covered with SouthernBiotech™ Fluoromount-G™ Slide Mounting Medium (Themo Fisher Scientific, cat #OB100-01) and a coverslip. Slides were kept in a light-protected box at 4°C before imaging by confocal microscopy.

### 2.5 Confocal Microscopy and Image Analysis

Immunostained tissue sections were imaged with an Olympus Fluoview FV300 confocal laser scanning microscope. Three regions of interest (ROI) in the proximal, middle, and distal thirds of one radial leaflet section per donor, encompassing the leaflet tip, belly, and base, were imaged at low magnification (10X) and mapped on the microscope stage. Within each ROI, z-stacks were acquired in sub-regions using a 40X objective. Confocal images were imported into FIJI software where maximum intensity z-projections of the Alexafluor488 (vimentin) and CY3 (α-SMA) channels were created. Otsu thresholding was used to segment the maximum intensity projections for each channel and quantify the positive signal area. For each ROI (tip, belly, base), the area of vimentin expression was summed across all sub-regions imaged at 40X. The same calculus was repeated to determine the total positive α-SMA area across the sub-regions that were sampled. The sum α-SMA positive area was calculated as a percentage of the sum vimentin positive area in each ROI.

### 2.6 Histology Staining, Imaging, and Analysis

To visualize ECM organization, leaflet sections serial to those used for immunostaining were used for Movat’s pentachrome staining (STARR Innovation Centre, University Health Network, Toronto, Canada). Stained sections were imaged at 40X magnification using a whole slide scanner (Aperio AT2 brightfield Scanner). Whole slide scans were viewed and analyzed in Aperio ImageScope Pathology Slide Viewing Software (Leica Biosystems). Three ROI with distinct lamellar structure in the belly of one radial leaflet section per donor were analyzed for thickness measurements of the fibrosa, spongiosa, and ventricularis layers. The outer margins of the leaflet section at the fibrosa and ventricularis surfaces, as well as those between each layer were annotated with free-form lines in each ROI. The distance measurement tool in ImageScope was used to calculate an average thickness measurement from 15 equidistantly placed measurements between the free form line annotations marking the boundaries of each layer. The thickness measurements for each layer were averaged across the sampled ROI to provide an average layer-specific thickness measurement for each PV donor.

### 2.7 Statistical Analysis and Data Visualization

For mechanical tests, statistical analyses were performed using JMP Pro 16 software. Data are reported as mean ± standard deviation (SEM). For pairwise comparison, data were analyzed using one-way ANOVA with Tukey’s multiple comparisons test. For quantification of ECM, DNA, and VIC phenotypes, statistical analyses were performed in GraphPad Prism 9 software using one-way ANOVA with post-hoc Tukey’s multiple comparisons test. Data are reported as mean ± standard error of mean (SEM). All data were visualized using GraphPad Prism 9.

## 3. Results

### 3.1 Biaxial Tensile Behaviour of Pediatric Pulmonary Valve Leaflet

Mechanical testing on fresh and frozen samples (N = 4) from matched donors showed that the snap-freezing process did not significantly impact the biaxial tension-strain behaviour (Figure 2A) and materials constants (p > 0.3; Figure 2B; Supplemental Table S2) of the pediatric PV tissue. Since testing fresh tissues posed a considerably greater logistical burden, biaxial mechanical testing on a larger number of donors (N = 9) was performed on tissues retrieved after one snap-freezing and thawing cycle. Pediatric PV tissues showed highly nonlinear, anisotropic biaxial tension-strain behaviour with larger extensibility in the radial direction of the valve (Figure 2C).

**Figure 2:**
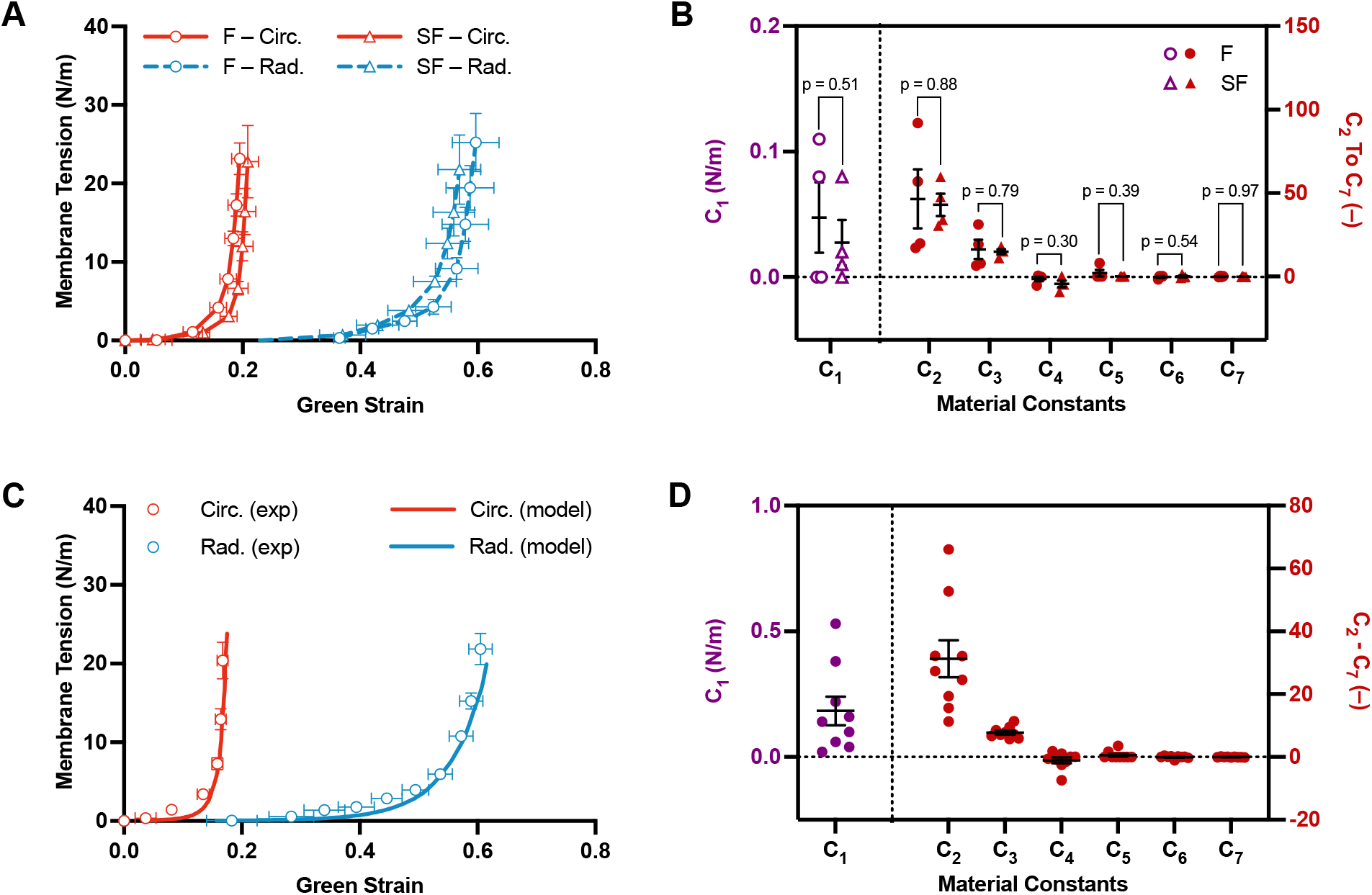
Tensile behaviour of piglet pulmonary valve (PV) under equibiaxial force. (A) Experimental biaxial tension-strain behaviour of fresh (F, triangle markers) and snap-frozen (SF, circle markers) PV tissue samples (N = 4), taken from matched donors, under equibiaxial loading (P5) in the circumferential (Circ., orange) and radial (Rad., blue) directions. (B) Material constants of fresh (F, purple and red triangle markers) and snap-frozen (SF, purple and red circle markers) PV tissue (N =4). (C) Experimental (exp) and model-fit (model) biaxial tension-strain behaviour of snap-frozen PV tissue (N = 9) in the circumferential (orange) and radial (blue) directions under equibiaxial loading (P5). (D) Material constants of snap-frozen PV tissues (N =9). All tests were performed at a strain rate of 0.08 s^-1^. Data represented as mean ± SEM, and scatter data points show biological replicates.

Among all material constants, C1, C2, and C3 were determined to be the most impactful in governing the mechanical behaviour of PV tissue (Figure 2D; Supplemental Table S2). Tension-strain behaviour of the tissue in the circumferential direction is mainly governed by C1 and C2 and that in the radial direction is governed by C1 and C3. Pearson correlation coefficients between the model and experimental data were 0.89 and 0.96 in the circumferential and radial directions, respectively, for all loading ratios (Supplemental Figure S2). For equibiaxial loading, the Pearson correlation coefficient was 0.99 in both circumferential and radial directions (Figure 2C).

### 3.2 Extracellular Matrix Composition

Three constituent ECM proteins of the native porcine PV (collagen, s-GAG, and elastin) were quantified by biochemical assays and normalized to either the wet weight of the sampled leaflet tissue or the lysate-matched DNA content (Figure 3). To determine if there was spatial heterogeneity in the DNA or tissue weight normalized ECM content of the leaflet across the regions sampled, results were stratified by the area of the leaflet from which the sample was taken. The spatial variability in the leaflet sampling was not reflected in either the weight or DNA-normalized hydroxyproline (Figure 3A, p = 0.28 and Figure 3I, p = 0.69) or s-GAG (Figure 3C, p = 0.94 and Figure 3K, p = 0.32) content of the PV. Similarly, the measured elastin content, which could only be normalized against the sampled tissue weight, as the DNA content of the oxalic assay lysates could not be quantified, did not change as a function of the spatial sampling (Figure 3E, p = 0.27). Finally, the total cellularity of the native leaflet based on the net DNA content measured was not different across the regions of the leaflet that were sampled (Figure 3G, p = 0.17).

**Figure 3:**
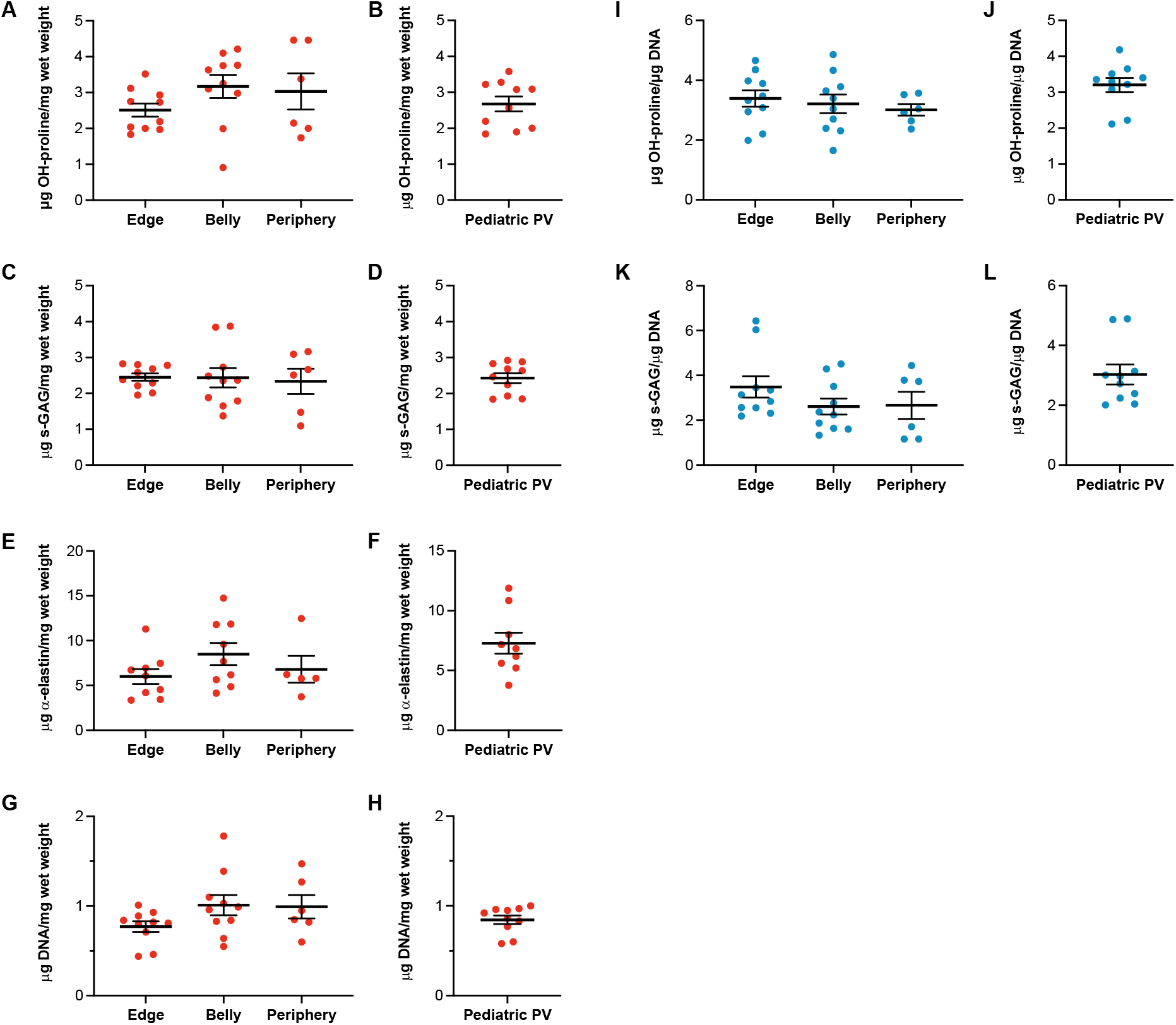
Tissue wet weight (Panels A-H, orange markers) and DNA normalized (Panels I-L, blue markers) quantification of piglet pulmonary valve (PV) extracellular matrix (ECM) with and without regional stratification. Tissue weight and DNA-normalized hydroxyproline (OH-proline) (A, I) and sulphated glycosaminoglycans (s-GAG) (C, K) content, and weight-normalized α-elastin (E) and DNA (G) did not vary based on the region of tissue sampling. The normalized sum across all sampled regions of the PV for all ECM proteins and DNA are shown in panels B, D, F, H, J, and L. Data represented as mean ± SEM and scatter plot shows biological replicates (N = 10 for edge and belly, N = 6 for periphery; N = 10 for whole PV).

Since regional heterogeneity was not reflected in either the ECM or net DNA content, we can provide benchmarks for PV leaflet tissue by taking a sum of the quantified ECM and DNA content across the edge, belly, and periphery of the leaflet that were sampled (Figure 3B, D, F, H, J, L and Table 1).

**Table 1:** Quantified extracellular matrix and DNA content in pediatric pulmonary valve (PV) with and without regional stratification. Values presented are mean ± SEM (N = 10). s-GAG: sulphated glycosminoglycans.

|  | Pediatric PV |  | Edge |  | Belly |  | Periphery |  |
| --- | --- | --- | --- | --- | --- | --- | --- | --- |
| *Normalization* | Weight<br>($\mu\text{g}/\text{mg}$) | DNA<br>($\mu\text{g}/\mu\text{g}$) | Weight<br>($\mu\text{g}/\text{mg}$) | DNA<br>($\mu\text{g}/\mu\text{g}$) | Weight<br>($\mu\text{g}/\text{mg}$) | DNA<br>($\mu\text{g}/\mu\text{g}$) | Weight<br>($\mu\text{g}/\text{mg}$) | DNA<br>($\text{g}/\mu\text{g}$) |
| **Hydroxyproline** | 2.68 ± 0.21 | 3.20 ± 0.20 | 2.51 ± 0.18 | 3.39 ± 0.28 | 3.17 ± 0.32 | 3.21 ± 0.31 | 3.03 ± 0.32 | 3.01 ± 0.19 |
| **s-GAG** | 2.43 ± 0.14 | 3.03 ± 0.33 | 2.45 ± 0.10 | 3.49 ± 0.48 | 2.43 ± 0.27 | 2.61 ± 0.36 | 2.33 ± 0.35 | 2.67 ± 0.61 |
| **α-Elastin** | 7.27 ± 0.88 | - | 6.00 ± 0.83 | - | 8.50 ± 1.23 | - | 6.80 ± 1.45 | - |
| **DNA** | 0.84 ± 0.05 | - | 0.77 ± 0.06 | - | 1.01 ± 0.11 | - | 0.99 ± 0.13 | - |

### 3.3 Extracellular Matrix Organization

Movat’s pentachrome staining of pediatric PV leaflet sections revealed a lamellar ECM organization with three layers: (i) the fibrosa populated by collagen, (ii) the spongiosa consisting of glycosaminoglycans, and (iii) the ventricularis composed of radially oriented elastin and collagen fibres (Figure 4B). On average, the native pediatric PV leaflets were 175.91 ± 12.23 μm thick, and the spongiosa layer accounted for the largest proportion of the leaflet thickness (95.99 ± 10.91 μm), followed by the fibrosa (57.89 ± 4.72 μm) and ventricularis (22.02 ± 2.07 μm) layers (Figure 4A).

**Figure 4:**
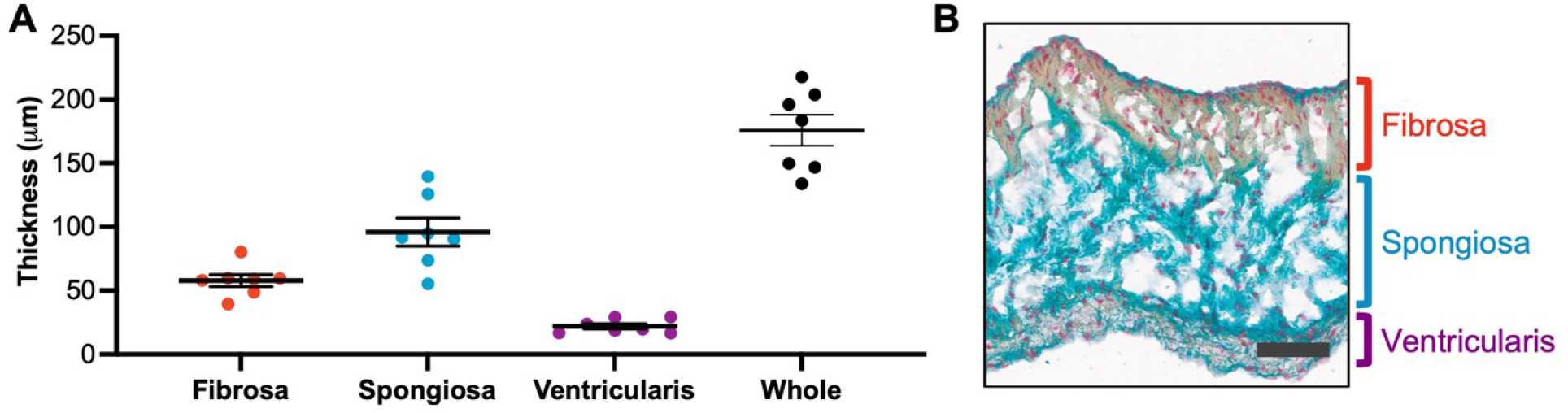
Extracellular matrix (ECM) organization and leaflet thickness in piglet pulmonary valve (PV) leaflet. (A) Whole leaflet and layer specific thickness of the native pediatric PV in the belly region. Data presented as mean ± SEM. Scatter plot shows biological replicates (N = 7). (B) Representative image of Movat’s pentachrome stained pediatric PV leaflet in the belly region showing a trilayer ECM organization. Red = nuclei, blue = proteoglycan, yellow = collagen, elastin = purple. Scale bar = 50 μm.

### 3.4 Valvular Interstitial Cell Phenotypes

Immunostaining of radial PV leaflet sections showed that the VIC population of the native pediatric leaflet primarily consisted of a quiescent vimentin-expressing fibroblast phenotype. α-SMA expression, indicative of a myofibroblast phenotype, increased moving down the radial axis of the leaflet, from the tip to the belly, and at the base (Figure 5A-C). Generally, α-SMA-expressing cells were sparsely distributed in the tip (Figure 5A) and belly (Figure 5B, top panel) of the leaflet. In regions with a comparatively higher proportion of α-SMA expressing VICs, such as parts of the belly (Figure 5B, bottom panel) and the base of the leaflet (Figure 5C), α-SMA expression was restricted to the ventricularis layer and surface as determined by Movat’s pentachrome staining of serial sections (Figure 5D). Quantitative analysis of α-SMA expression in the leaflet demonstrated that, on average, α-SMA positive area as a percentage of the vimentin positive area was 0.50 ± 0.05% in the tip, 2.47 ± 1.21% in the belly, and 4.17 ± 1.15% in the base of the leaflet (mean ± SEM), with a statistically significant difference between the tip and base of the leaflet (p = 0.029) (Figure 5E).

**Figure 5:**
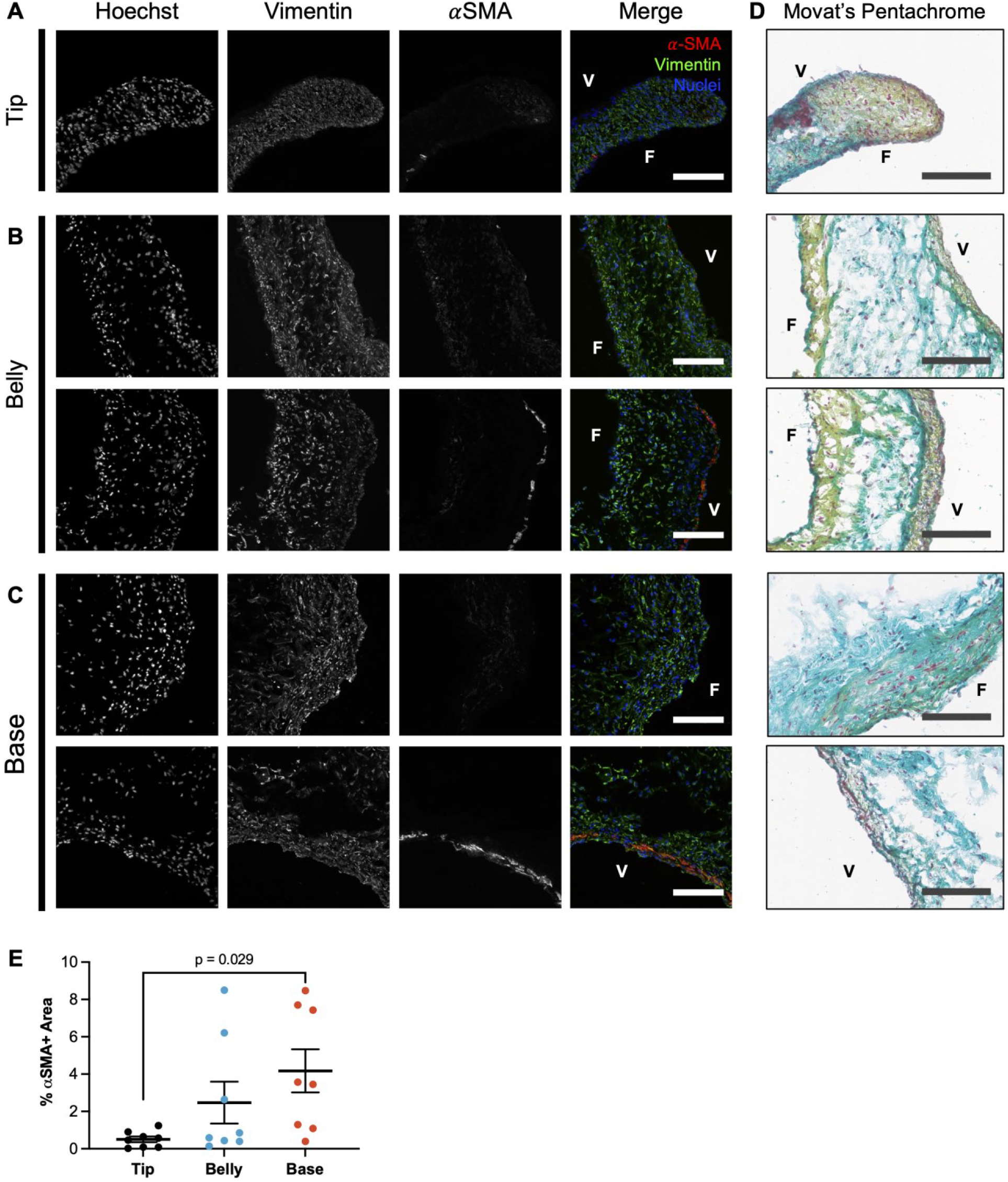
**α**-smooth muscle actin (α-SMA) and vimentin expression in native piglet pulmonary valve (PV) leaflet. (A-C) Representative images of α-SMA and vimentin expression in the tip, belly, and base of the PV leaflet. Valvular interstitial cells (VICs) sparsely expressed α-SMA in the tip of the leaflet (A). α-SMA expression in the belly of the leaflet was more variable, with representative images reflecting regions of minimal (B, top panel) and maximal (B, bottom panel) expression. The base of the leaflet consisted of the highest proportion of α-SMA expressing VICs, with representative images showing the fibrosa (C, top panel) and ventricularis (C, bottom panel) surfaces of the leaflet. Comparison of immunostained PV tissue against their Movat’s pentachrome stained serial sections (D) showed that α-SMA expression was restricted to the ventricularis surface of the leaflet in the belly and base. (E) Quantitative analysis of immunostained tissue sections demonstrated that α-SMA positive area (normalized to vimentin positive area) increased moving from the tip to the belly and base of the leaflet. Data presented as mean ± SEM. Scatter plot shows biological replicates (N = 8). Scale bars = 100 μm. (V: ventricularis; F: fibrosa)

## 4. Discussion

While many HVTE strategies are motivated by a clinical need for living PV tissue capable of somatic growth and remodeling in a pediatric recipient, few have evaluated bioengineered constructs against native pediatric PV tissue. This is due to the sparse and incomplete understanding of the relevant properties of the human pediatric PV, stemming from limited access to healthy pediatric tissues. The need to address this knowledge gap is underscored by the lack of long-term functionality among TEHVs produced to date, which suggests that bioengineered constructs should better approximate the native tissue prior to implantation. In the present study, we used the piglet PV as a model for the native human pediatric PV and characterized the planar biaxial tensile behaviour, ECM composition and organization, and VIC phenotypes across 10 donor valves.

To the best of our knowledge, this study is the first to conduct planar biaxial tensile testing of pediatric PV tissue, independent of donor species, from which a constitutive model of the native pediatric leaflet was derived. Typically, the mechanical behaviour of TEHVs and pediatric PV and TEHVs have been compared on the basis of uniaxial tensile tests of circumferential and radial tissue strips^35-37^. While uniaxial tests of anisotropic tissues like the native valve yield a non-linear stress-strain profile, they involve non-physiologic loading protocols where fibre kinematics, namely the straightening of circumferentially oriented crimped collagen fibres and the rotation of these fibres towards the stretch axis, are not preserved^38^. Further, most studies rely on a single metric, the tangential elastic modulus, to characterize the mechanical behavior of the native valve. However, the tangential modulus provides only a snapshot into the complex non-linear tensile behaviour of the native leaflet in the circumferential and radial axes. By contrast, a predictive constitutive model enables recapitulation of the tension-strain profile in its entirety and under various loading conditions beyond those that have been experimentally tested. In fact, in a previous study of the biaxial planar tensile behaviour of native pediatric PV from two donor tissues (8 months and 11 years old)^15^, the lack of constitutive modelling for complete description and prediction of mechanical behaviour of the leaflets was acknowledged as a limitation of the study. In the same study, authors raised the need to address the effects of cryopreservation on biomechanics of the valve. The need for a constitutive tensile model is especially pertinent to recent work focused on creating mechanically biomimetic constructs that fully recapitulate the leaflet’s stress-strain response^39-41^.

In this study, we have addressed all of the aforementioned gaps. We used a planar biaxial tensile testing protocol^30^ to test the native piglet PV under equibiaxial and various proportional loading protocols. This detailed dataset was subsequently analyzed to derive material constants for a constitutive seven-parameter strain energy function (Fung Model) that can be used to describe the tensile behaviour of native piglet PV leaflet in the radial and circumferential axes under various loading conditions. The Pearson correlation coefficients of 0.89 and 0.96 calculated in the model fit test for the circumferential and radial axes, respectively, reflected the overall match between the experimental and predicted data for all loading protocols. Ultimately, we found that the native pediatric PV has a highly non-linear, anisotropic biaxial tension-strain response, which aligns with previous literature characterizing other semilunar valves^30,31,42-45^. Furthermore, by conducting tests on fresh and donor-matched snap-frozen leaflets of four donors, we demonstrated that the tensile behaviour of the native leaflet is preserved after one freeze-thawing cycle. These findings motivated subsequent tests on cryopreserved tissues, which removed the logistical burden of processing fresh tissues for characterization studies.

In this study we also quantified the constituent ECM proteins of the native piglet PV, which were normalized against the wet weight and DNA content of the samples. We found that stratifying the quantified ECM content based on sampling region did not reveal any significant differences in normalized ECM content (DNA and wet weight normalized) or cellularity, suggesting homogenous distribution of tissue and cells in the pediatric PV leaflet in the circumferential-radial planar axis. These findings are consistent with reports of relatively even distribution of collagen fibres and smaller fibre bundles observed in pediatric human PV (at 2 months and 2 years-old) than in the adult equivalent^23^. However, it is possible that the sample size from each designated region in this study was too large to provide the spatial resolution necessary to detect differences in regional ECM and cell content. The absence of spatial heterogeneity reported here allowed us to present the sum of the ECM content across all sampled regions normalized to the sum of the tissue wet weight or DNA from the same regions to provide an estimate of the ECM content of the piglet PV as a whole. While previous studies have quantified the ECM content of pediatric human PV^15^ and piglet PV^48^, the results are presented as normalized to dry weight of lyophilized leaflet tissue, making it difficult to draw direct comparison to the quantified ECM values as reported in this study. However, as expected, the dry weight normalized hydroxyproline and s-GAG content of the pediatric human and porcine PVs are greater than the wet weight normalized ECM quantified here. This is attributable to the fact that dry weight normalization does not include the water weight retained in the spongiosa of the leaflet, and thus, quantified ECM content is normalized to a smaller mass.

Our histochemical stains of the piglet PV revealed a trilayer construct with distinct fibrosa, spongiosa, and ventricularis layers. This finding is consistent with previous studies of ECM architecture in the human pediatric PV^15,16^ and piglet PV^48^. The significant contribution of the spongiosa layer to the thickness of the native pediatric PV leaflet has also been qualitatively observed in previous studies, and is representative of the transition between the fetal valve which is primarily composed of GAGs and the adult valve consisting primarily of dense collagen and elastin^15,16^. In addition to visualizing the native ECM organization, we used Movat’s pentachrome stained piglet PV leaflets to quantify layer specific thickness of the leaflet. These measurements provide thickness benchmarks for evaluation of TEHV constructs in both studies of whole TEHV leaflets and layer-specific constructs ^49-51^.

Cell-mediated contraction of TEHVs is a consistent limitation reported in the literature to date^52^. This is thought to be a function of a disproportionate population of activated α-SMA expressing interstitial cells. While activated myofibroblastic cells are necessary to synthesize neotissue, unrestrained activation may be the cause of leaflet dysfunction in bioengineered valves. Quantifying the proportion of α-SMA expressing VICs in the native piglet PV can provide benchmarks for a bioengineered pediatric construct and for TEHVs as a whole, since they provide a snapshot into the cell population at a stage of rapid somatic growth and development. Our work has demonstrated that α-SMA expression, quantified as α-SMA positive area as a proportion of vimentin positive area, followed a pattern of increasing expression from the tip to the base of the leaflet. In regions of maximal expression, the α-SMA positive area accounted for less than 5% of the vimentin positive area. The immunostaining also revealed that α-SMA-expressing cells were restricted to the subendothelial layer on the ventricularis surface of the leaflet. Our findings are partially consistent with those of Aikawa *et al*.^16^ who previously reported that α-SMA expressing cells accounted for, on average, 6.2% of VICs in the human pediatric PV leaflet (6.0 ± 1.6 y.o) and resided in the subendothelial layer of the outflow (fibrosa) surface of the leaflet. The partial corroboration between the results reported here and those of previous studies on the human pediatric PV with respect to ECM organization and VIC phenotypes suggests that the piglet PV model is an appropriate analogue for the pediatric human tissue in this context.

Though adult PV leaflets were not evaluated in this study, previous findings of age-related changes in the structure and mechanical behaviour of heart valves suggest that the pediatric tissue has unique properties that are not matched by its adult counterpart, thus requiring independent design goals for a bioengineered replacement. Age-dependent trends in cellular phenotypes, ECM composition, and/or mechanical behaviour have been reported in human^15-17,23,24^, ovine^53^, and porcine^54-57^ semilunar valves, although only a fraction of these studies include the PV in their analysis^15,16,23,53^. In a seminal study, Aikawa *et al*. reported on age-related changes in ECM content and organization, and VIC phenotypes of the PV, namely: (i) maturation of the lamellar ECM organization evidenced by increasing collagen fibre thickness, elastin content, and uniformity of collagen fibre alignment as a function of age; and (ii) a shift from an activated myofibroblast phenotype in pediatric PV VICs to a quiescent fibroblast phenotype in their adult counterpart. van Geemen *et al.* found patterns of increasing tangential stiffness (at both low and high strains) and collagen cross-link density in parallel with decreasing radial and circumferential extensibility, s-GAG content, cellularity, and proportion of α-SMA expressing VICs between pediatric and adult human PVs^15^. Corroborating the findings of both studies, Oomen et al. reported that circumferentially oriented collagen fibres organized into larger collagen fibre bundles in the aging PV ^23^.

With the exception of the work by van Geemen *et al.*, studies of age-related changes in the mechanical behaviour of the native semilunar valves have focused on the aortic valve. Christie and Barratt-Boyes found that radial extensibility of the human aortic valve leaflet in biaxial tensile tests was negatively correlated with donor age, with the first and most rapid decline occurring in late adolescence^24^. This trend was later observed in porcine aortic valves under uniaxial tensile testing^56^. Whether the same changes occur in the aging PV is less clear given the limited data on the tensile behaviour of the pediatric PV. Moreover, reported differences between the two semilunar valves challenge the extrapolation of age-related changes in the structure and function of the aortic valve to the PV. In the adult population, displacement-controlled uniaxial tensile tests of the human^58^ and porcine^59^ semilunar valves have demonstrated that the ultimate stress in the circumferential axis and ultimate strain in the radial axis of the PV are significantly greater than those of the aortic valve^58^. These findings are corroborated by physiological force-controlled biaxial tensile tests, which demonstrated that circumferential strains in the porcine PV are lower than those in the aortic valve with an acute transition point following the toe region under equibiaxial tensile loads^60^, likely due to the PV’s comparatively smaller proportion of crimped collagen fibres, smaller collagen crimp amplitude, and greater alignment of collagen fibres in the unloaded state^61^.

It is possible that the reported differences between the PV and aortic valve are unique to the adult population, and the properties of the pediatric aortic valve could serve as design surrogates for the pediatric PV. Existing literature comparing the PV and aortic valve at different stages of development and aging suggests that the differences between the semilunar valves are either transient or only become apparent in adolescence and/or adulthood. For instance, Aikawa *et al*. found that the proportion of activated α-SMA expressing VICs was significantly higher in the aortic valve than PV only during the late neonatal period (<30 days-old), owing to the rapid decline in VIC activation levels in the PV between early (minutes to hours old) and late neonatal life^16^. This was attributed to the functional response of VICs to the changing hemodynamic conditions during the transition from fetal to postnatal circulation, wherein pulmonary pressures decrease (30/15 mmHg) and aortic pressures increase (70/40 mmHg) after both being exposed to equal arterial pressures (50/15 mmHg) in utero^16^. By childhood, VIC activation levels were identical in the healthy PV and aortic valve^16^. Significant differences in leaflet thickness between the PV and aortic valve are evident in adolescence and adulthood^15,58,59^, potentially due to the increased elastin content and ventricularis thickness of the adult aortic valve versus adult PV^16^. Lending further credence to the argument that the pediatric PV and aortic valve have interchangeable properties are similarities in the weight-normalized collagen, s-GAG and DNA content of the two valves ^15^. However, measurements of bulk ECM content and lamellar organization do not offer insight into potential differences in the microarchitecture of collagen fibres in the pediatric semilunar valves, namely the density of crimped collagen fibres, crimp period and amplitude, and orientation, which account for the reported differences in the tensile behaviour of the adult aortic valve and PV^60-62^. The current body of literature cannot address whether these microarchitectural differences also exist between the pediatric semilunar valves, presumably due to the aforementioned challenges in accessing healthy pediatric valves, suggesting that the most accurate benchmark for bioengineered pediatric PV tissue is the native pediatric PV and not its left-sided counterpart.

## 5. Conclusion

The native piglet PV leaflet is an anisotropic construct with a non-linear tension-strain behavior in both the radial and circumferential directions. The pediatric leaflet is stratified into three distinct ECM layers – fibrosa, spongiosa, and ventricularis – wherein the spongiosa accounts for a majority of the leaflet thickness. The VIC population in the piglet PV, consists of a small proportion of activated α-SMA expressing cells located primarily at the base of the leaflet and on the ventricularis surface. Together, our findings provide standards to guide the development of tissue engineered pediatric PVs.

## Notes

### Competing Interest Statement

The authors have declared no competing interest.

