## Supplemental Information for "Characterization of pediatric porcine pulmonary valves as a model for tissue engineered heart valves"

### Characterization of pediatric porcine pulmonary valve as a model for the tissue engineered pediatric valve


#### Supplemental Materials and Methods

##### Processing of Piglet Pulmonary Valve Tissue

Three pulmonary valve (PV) leaflets from 6 donors were used in this study (one fresh and two snap-frozen leaflets), while only two snap-frozen leaflets were analyzed from 4 donors. Leaflets were processed for biaxial tensile testing, biochemical analysis, immunostaining and histology, as described in Figure 1. In the first snap-frozen leaflet, a parallel cutter was used to isolate a radial strip (4.5 mm wide) spanning the length of the leaflet from the basal attachment to the free margin (Figure 1, (1)). The remainder of the leaflet on either side, encompassing tissue between either commissure and the belly, were denoted as the leaflet edge, and subsequently digested in either papain or oxalic acid. A second cut (Figure 1, (2)), in the circumferential axis of the leaflet, was made to the radial strip to isolate a square sample from the belly of the leaflet for tensile testing. The free margin and basal attachment regions of the radial strip were stored in phosphate-buffered saline (PBS) during tensile testing of the belly region (yellow square). Following mechanical testing, the square sample from the belly was rinsed in PBS to remove graphite particles. A third cut (Figure 1, (3)) was then made in the circumferential axis of the square sample. The top half of the square sample was combined with the free margin remnant from the second cut and digested in papain. The bottom half of the square sample was combined with the basal attachment remnant from the second cut and digested in oxalic acid (OA). Both lysates were given the spatial designation of the belly region. The second leaflet was cut along the radial axis at the midline of the leaflet (Figure 1, (4)). One half of the leaflet was embedded in optimal cutting temperature (OCT) compound and snap-frozen using liquid nitrogen and stored at -80 °C before cryosectioning. From the remaining half of the second leaflet, three biopsy punches (2 mm diameter) were taken for an unrelated study. The remaining peripheral tissue of the leaflet was divided in half (Figure 1, (5)) and digested in either papain or OA.

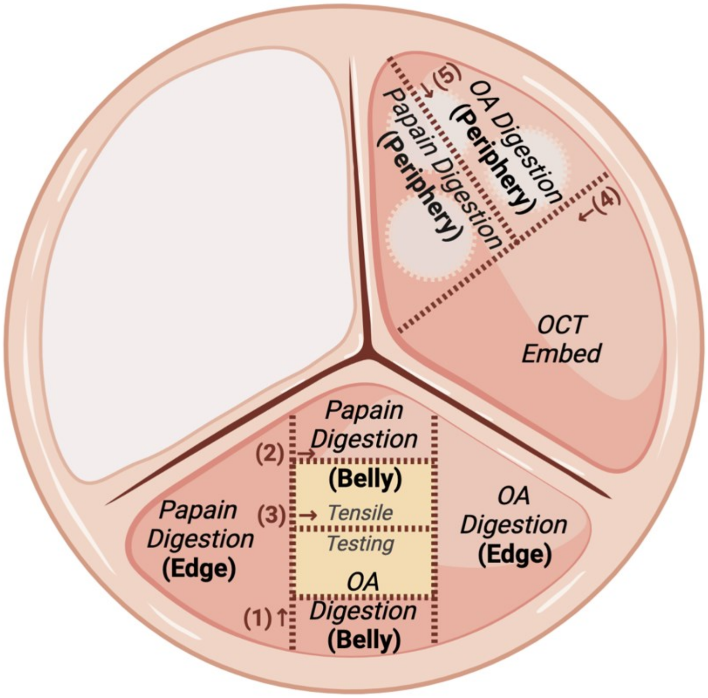


**Figure S1:** Overview of pulmonary valve (PV) leaflet tissue processing for characterization of mechanical behaviour, extracellular matrix composition and organization, and valvular interstitial cell phenotypes. Gray shading indicates regions or whole leaflets that were tested fresh or used in unrelated studies. Dashed lines indicates where leaflet tissue was cut. OA: oxalic acid; OCT: optimal cutting temperature.

**Equibiaxial Force Displacement-Controlled Testing**

Tissues were subjected to ten displacement-controlled preconditioning cycles with displacements of $U_{Max,Circ.}$ and $U_{Max,Rad.}$ in the circumferential and radial direction, respectively, that generated equibiaxial loading in the tissue. Subsequently, they underwent nine displacement-controlled testing protocols with gradually increasing circumferential-to-radial load ratios as described in Table 1.

**Table S1:** Prescribed displacement in the circumferential and radial direction of pulmonary valve tissue samples in the preconditioning and test cycles.

| **Cycles** | **Displacement in the circumferential direction** | **Displacement in the radial direction** |
|---|---|---|
| Preconditioning | $U_{Max,Circ.}$ | $U_{Max,Rad.}$ |
| Test cycle 1 (P1) | $U_{Max,Circ.} + U_{Adj,Circ.}$ | $\frac{6}{10} U_{Max,Rad.}$ |
| Test cycle 2 (P2) | $U_{Max,Circ.} + \frac{3}{4} U_{Adj,Circ.}$ | $\frac{7}{10} U_{Max,Rad.}$ |
| Test cycle 3 (P3) | $U_{Max,Circ.} + \frac{1}{2} U_{Adj,Circ.}$ | $\frac{8}{10} U_{Max,Rad.}$ |
| Test cycle 4 (P4) | $U_{Max,Circ.} + \frac{1}{4} U_{Adj,Circ.}$ | $\frac{9}{10} U_{Max,Rad.}$ |
| Test cycle 5 (P5) | $U_{Max,Circ.}$ | $U_{Max,Rad.}$ |
| Test cycle 6 (P6) | $\frac{4}{5} U_{Max,Circ.}$ | $U_{Max,Rad.} + \frac{1}{4} U_{Adj,Rad.}$ |
| Test cycle 7 (P7) | $\frac{3}{5} U_{Max,Circ.}$ | $U_{Max,Rad.} + \frac{1}{2} U_{Adj,Rad.}$ |
| Test cycle 8 (P8) | $\frac{2}{5} U_{Max,Circ.}$ | $U_{Max,Rad.} + \frac{3}{4} U_{Adj,Rad.}$ |
| Test cycle 9 (P9) | $\frac{1}{5} U_{Max,Circ.}$ | $U_{Max,Rad.} + U_{Adj,Rad.}$ |

#### Analysis of Biaxial Tensile Testing Data

The method used for the calculation of Green strain, membrane tension, and material constants was reported in detail previously [1]. In brief, the original domain in the sample is defined as $-L/2 \leq X_1 \leq L/2$, $-L/2 \leq X_2 \leq L/2$, and $-H/2 \leq X_3 \leq H/2$ in the sample's undeformed state, where $L$ is the in-plane width and length of the tissue sample; H is its thickness; and $X_1$, $X_2$, and $X_3$ correspond to the X, Y, and Z coordinates, respectively. During the nine test protocols (P1–P9), the location of the tracked points, determined by image analysis, are mapped to $x_1$, $x_2$, and $x_3$ using $x_1 = \lambda_1 X_1 + F_{12} X_2$, $x_2 = F_{21} X_1 + \lambda_2 X_2$, and $x_3 = (h/H)\, X_3$, where $F_{12}$ and $F_{21}$ are the components of the transformation gradient tensor:

$$\boldsymbol{F} = \partial \boldsymbol{x} / \partial \boldsymbol{X} = \begin{bmatrix} \lambda_1 & F_{12} & 0 \\ F_{21} & \lambda_2 & 0 \\ 0 & 0 & h/H \end{bmatrix} \quad (1)$$

The thickness of the sample during deformation can be calculated using $h = H/J_{2D}$, where $J_{2D} = \lambda_1 \lambda_2 - F_{12} F_{21}$, based on the assumption that the determinant of the transformation gradient tensor is equal to 1, as the tissue is approximately incompressible.

The Green strain tensor can be calculated using $\boldsymbol{E} = 1/2\,(\boldsymbol{F}^T \cdot \boldsymbol{F} - \boldsymbol{I})$, where the dot represents the scalar product of two tensors. The components of the membrane tension tensor, $\boldsymbol{S}$, were calculated according to:

$$T_{11}^{S} = \lambda_2 f_1 / (J_{2D} L) \quad (2)$$

$$T_{22}^{S} = \lambda_1 f_2/(J_{2D}L) \quad (3)$$

where $f_1$ and $f_2$ are the force readings in the X and Y directions, respectively. The resultant $T_{11}^{S}$-$E_{11}$ and $T_{22}^{S}$-$E_{22}$ curves were plotted to represent the experimental membrane tension-Green strain relationships in X and Y direction of the PV tissue.

Assuming that the strain energy is conserved in the test cycles (P1–P9), the tension and Green strain tensors are related according to:

$$\begin{bmatrix} T_{11}^{S} & T_{12}^{S} \\ T_{12}^{S} & T_{22}^{S} \end{bmatrix} = \begin{bmatrix} \frac{\partial w}{\partial E_{11}} & \frac{\partial w}{\partial E_{12}} \\ \frac{\partial w}{\partial E_{12}} & \frac{\partial w}{\partial E_{22}} \end{bmatrix} \quad (4)$$

where $w$ is the two-dimensonal strain energy function.

To characterize the mechanical properties of the PV tissue via material constants, a seven-parameter Fung model [1] was used:

$$w = \frac{C_1}{2}(e^{Q} - 1) \quad (5)$$

where $Q = C_2{E_{11}}^2 + C_3{E_{22}}^2 + 2C_4E_{11}E_{22} + C_5{E_{12}}^2 + 2C_6E_{11}E_{12} + 2C_7E_{22}E_{12}$.

Material constants ($C_1$– $C_7$) were determined by minimizing the objective function, $g$, matching the experimental and model-predicted membrane tensions:

$$g = \left\| \left|T_{11_{Exp}}^{S} - T_{11_{Model}}^{S}\right| + \left|T_{22_{Exp}}^{S} - T_{22_{Model}}^{S}\right| + \left|T_{12_{Exp}}^{S} - T_{12_{Model}}^{S}\right| \right\|^2 \quad (6)$$

where

$T_{11_{Exp}}^{S} = \lambda_2 f_1/(J_{2D}L)$ and $T_{11_{Model}}^{S} = C_1e^{Q}(C_2E_{11} + C_4E_{22} + C_6E_{12}$;
$T_{22_{Exp}}^{S} = \lambda_1 f_2/(J_{2D}L)$ and $T_{11_{Model}}^{S} = C_1e^{Q}(C_3E_{22} + C_4E_{11} + C_7E_{12}$;
$T_{12_{Exp}}^{S} = F_{21} f_1/(J_{2D}L)$ and $T_{12_{Model}}^{S} = C_1e^{Q}(C_5E_{12} + C_6E_{11} + C_7E_{22})$.

All calculations were performed using a previously developed MATLAB code [1].

**DNA Quantification**

Total DNA content of papain lysates was quantified using the Hoechst dye 33258 assay [2]. Papain digested samples were thawed on ice and 50 μL of concentrated or diluted lysate (1:3-1:5 dilution depending on wet weight of sample) was aliquoted into a black wall, clear bottom 96-well plate in 2-4 replicates. Samples were mixed with 200 μL 0.1 μg/mL Hoechst 33258 dye (Life Technologies, cat. #H3569). The well plate was then transferred on ice to the plate reader and sample fluorescence was measured (excitation at 350 nm and emission at 450 nm). The net DNA content of the samples was quantified against calf thymus DNA (Sigma, cat. #D3664) standards of known concentrations.

**Sulphated Glycosaminoglycan Quantification**

Papain digested samples were thawed on ice and 20 μL of lysate was aliquoted into a 96-well plate in triplicate. Samples were mixed with 200 μL 0.016% dimethylmethylene blue dye (Sigma, cat. #341088) solution. The well plate was transferred to the plate reader and absorbance was measured at 525 nm. Sulphated glycosaminoglycan content was quantified against chondroitin sulfate standards (Sigma, cat. #C9819) of known concentrations.

##### Hydroxyproline Quantification

Total hydroxyproline content of papain lysates was quantified using the chloramine-T/Ehrlich's reagent assay [3]. Papain digested samples were thawed on ice and 100 μL of lysate was mixed with 100 μL 6.0 N HCl in heat-resistant glass culture tubes with screw caps. The mixture was heated at 110°C on a heat-block for 18 hours. The hydroxylates were taken off the heat-block and neutralized with 100 μL 5.7 N NaOH. The neutralized samples were further diluted with deionized water. Final dilution of papain lysates, including HCl, NaOH and deionized water volumes, ranged from 1:4-1:8 depending on the wet weight of the sample. 100 μL of neutralized samples were aliquoted into a 96-well plate in triplicate. Aliquots were subsequently mixed with 50 μL 0.05 N chloramine-T (Sigma, cat. #857319) (20 minutes, room temperature), followed by 50 μL 3.15 N perchloric acid (5 minutes, room temperature) and finally 50 μL Ehrlich's reagent (Sigma, cat. #39070) (20 minutes, 60°C). The well plate was cooled on ice and transferred to the plate reader. Absorbance was measured at 560 nm. Hydroxyproline content was measured against L-hydroxyproline (Sigma, cat. #H54409) standard solutions of known concentration.

##### Insoluble α-elastin Quantification

Total insoluble elastin content was measured using the Fastin™ Elastin Assay Kit by Biocolor (Accurate Chemical and Scientific Corporation, cat. # CLRF2000). The full volume of the oxalic acid lysate (750 μL) was used as a single replicate to quantify insoluble α-elastin content following the manufacturer's protocol.

#### Supplemental Results

**Table S2:** Experimentally derived material constants for 7-parameter Fung Model for snap-frozen pediatric porcine pulmonary valve (PV) leaflet tissue. Data presented as mean ± SEM (N = 9).

| Constant | Snap-Frozen Pediatric PV |
|---|---|
| $C_1$ | 0.18 ± 0.06 |
| $C_2$ | 31.25 ± 5.91 |
| $C_3$ | 7.77 ± 0.60 |
| $C_4$ | -1.07 ± 0.90 |
| $C_5$ | 0.59 ± 0.42 |
| $C_6$ | -0.04 ± 0.14 |
| $C_7$ | -0.01 ± 0.03 |

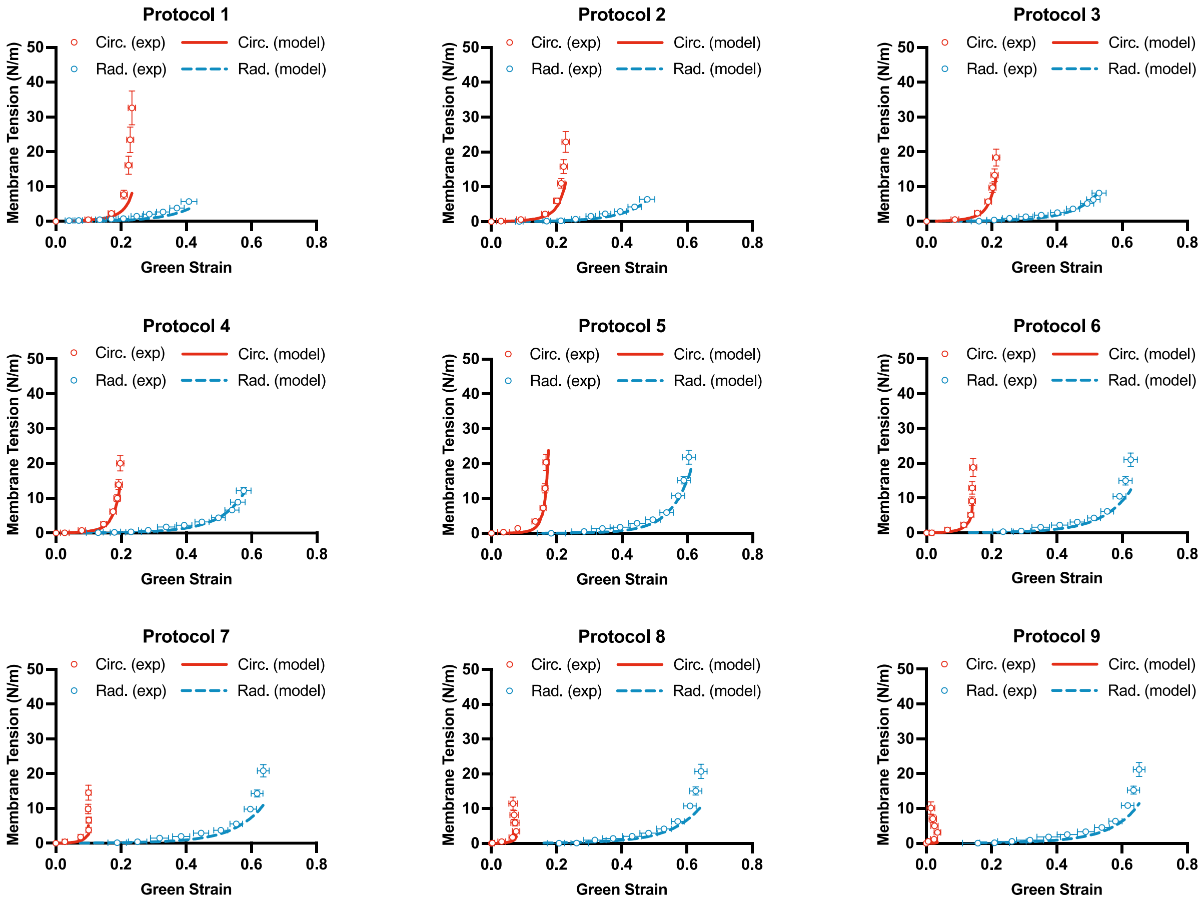


**Figure S2:** Tensile behaviour of piglet pulmonary valve (PV) under proportional biaxial loads described in Table 1. The experimental (exp) and model-predicted (model) biaxial tension-strain behaviour of snap-frozen PV samples in nine test cycle protocols (P1–P9). Data represented as mean ± SEM (N = 9).
